# *In situ* abundance and carbon fixation activity of distinct anoxygenic phototrophs in the stratified seawater lake Rogoznica

**DOI:** 10.1101/631366

**Authors:** Petra Pjevac, Stefan Dyksma, Tobias Goldhammer, Izabela Mujakić, Michal Koblížek, Marc Mussmann, Rudolf Amann, Sandi Orlić

## Abstract

Sulfide-driven anoxygenic photosynthesis is an ancient microbial metabolism that contributes significantly to inorganic carbon fixation in stratified, sulfidic water bodies. Methods commonly applied to quantify inorganic carbon fixation by anoxygenic phototrophs, however, cannot resolve the contributions of distinct microbial populations to the overall process. We implemented a straightforward workflow, consisting of radioisotope labeling and flow cytometric cell sorting based on the distinct autofluorescence of bacterial photo pigments, to discriminate and quantify contributions of co-occurring anoxygenic phototrophic populations to *in situ* inorganic carbon fixation in environmental samples. This allowed us to assign 89.3 ±7.6% of daytime inorganic carbon fixation by anoxygenic phototrophs in Lake Rogoznica (Croatia) to an abundant chemocline-dwelling population of green sulfur bacteria (dominated by *Chlorobium phaeobacteroides*), whereas the co-occurring purple sulfur bacteria (*Halochromatium* sp.) contributed only 1.8 ±1.4%. Furthermore, we obtained two metagenome assembled genomes of green sulfur bacteria and one of a purple sulfur bacterium which provides the first genomic insights into the genus *Halochromatium*, confirming its high metabolic flexibility and physiological potential for mixo-and heterotrophic growth.

## Introduction

Modern anoxic phototrophic ecosystems are often seen as analogs to study the ecology of early Earth environments. Sulfide-driven, anoxygenic photosynthesis is an ancient bacterial energy-yielding metabolism (Brocks et al., 2005) and often dominates autotrophic carbon fixation in stratified, sulfidic environments (e.g. Cohen et al., 1977). This process is primarily mediated by two phylogenetically distinct groups of bacteria: i) the strictly anaerobic green sulfur bacteria (GSB) of the class *Chlorobia* (Overmann, 2006), and ii) the metabolically more versatile purple sulfur bacteria (PSB) affiliated with the gammaproteobacterial orders *Chromatiales* and *Ectothiorhodospirales* (Imhoff, 2006a, 2006b). GSB and PSB oxidize and thereby detoxify sulfide and other reduced sulfur compounds, primarily formed during sulfate-dependent mineralization of organic matter in anoxic waters and sediments of marine and limnic environments (Overmann and Garcia-Pichel, 2013). Although GSB and PSB basically compete for the same resources, they co-occur in most photic and sulfidic environments. Ecological niche partitioning between GSB and PSB is considered to be mainly based on sulfide and oxygen concentrations, and light availability (Abella et al., 1980; Mas and van Gemerden, 1995; Stomp et al., 2007). PSB require more light and typically form dense populations at the sulfide-oxygen interface, while low-light adapted GSB often thrive beneath the PSB layer (e.g. Musat et al., 2008; Llorens-Marès et al., 2015). While GSB are strictly photoautotrophic, some PSB are capable of mixo-or heterotrophic, and chemotrophic growth (e.g. Imhoff, 2006a, 2006b; Berg et al., 2019), which further facilitates niche partitioning.

Numerous studies have addressed the overall contribution of anoxygenic photosynthesis (by GSB and PSB) to *in situ* dissolved inorganic carbon (DIC) fixation in sulfidic, stratified aquatic environments (e.g. Camacho et al., 2001; Casamayor et al., 2001; Marschall et al., 2010; Fontes et al., 2011; Morana et al., 2016). However, only few studies have investigated the relative contributions of GSB and PSB populations to this process. Musat and colleagues (2008) applied <u>nano</u>scale <u>s</u>econdary ion <u>m</u>ass <u>s</u>pectrometry (nanoSIMS) to track uptake of ^13^C-bicarbonate and ^15^ N-ammonia in single cells of GSB and PSB from the chemocline of Lake Cadagno. They reported that relative abundances of PSB and GSB do not correlate with their DIC fixation activity. Instead, the contribution of a rare PSB population (*Chromatium okenii*) to total DIC fixation was disproportionally high. In fact, per cell DIC fixation rates in *Chr. okenii* were up to three orders of magnitude higher than per cell DIC fixation rates of the *in situ* dominant GSB *Chlorobium clathratiforme*. In another study at Lake Cadagno, Storelli and colleagues (2013) incubated pure cultures of PSB and GSB isolated from Lake Cadagno *in situ* with ^14^C-labeled bicarbonate and quantified their contribution to DIC fixation by scintillographic quantification of radioisotope incorporation into microbial biomass. Again, a disproportionately high rate of DIC fixation in cells of *in situ* rare PSBs (*Candidatus* Thiocystis syntrophicum and *Lamprocystis purpurea*) was observed, while DIC fixation by the *in situ* dominant GSB *Chl. clathratiforme* was only marginal. However, these studies were somewhat limited by incubations performed *ex situ* or use of pure cultures that did not necessarily represent the *in situ* PSB and GSB communities. Thus, it is still unknown whether rare populations of PSB in general account for high proportions of DIC fixation via anoxygenic photosynthesis, or if this phenomenon is specific to Lake Cadagno.

Here, we present an approach combining incubations with radioisotope-labeled bicarbonate, fluorescence activated cell sorting (FACS) and scintillography of the sorted populations to discriminate the contribution of environmental populations of PSB and GSB to *in situ* DIC fixation. Similar approaches have previously been used to quantify the activities of discrete microbial populations in both pelagic and benthic environments (e.g. Zubkov et al., 2003; Vila-Costa et al., 2006; Dyksma et al., 2016). These high throughput workflows enable researches to directly measure the relative contribution of individual microbial groups to an overall process, without relying on assumptions such as conversion factors. Using this approach, we were able to show that the *in situ* numerically dominant GSB also dominate DIC fixation in our model system - a stratified seawater lake (Lake Rogoznica) on the eastern Adriatic coast (Croatia). The *in situ* rare PSB, on the other hand, contributed only marginally to DIC fixation in Lake Rogoznica. The metagenome assembled genome of these PSB, representing the first genomic dataset for the genus *Halochromatium*, illustrates their capability of mixo-and heterotrophic growth.

## Results and Discussion

### Chemical and hydrographical settings

Lake Rogoznica is a monomictic (stratified, except for one annual mixing) seawater lake on the eastern Adriatic coast (Fig. S1). At the time of sampling and experimentation (April, 2015), the water column of Lake Rogoznica was stratified. The recorded chemical and hydrographical water profiles were in accordance with data collected on numerous occasions in spring time over the last three decades (e.g. Bura-Nakić et al., 2009; Pjevac et al., 2015). The salinity of the deeper water layers was similar to the surrounding Adriatic seawater source (Lipizer et al., 2014). The decreased salinity in the top 4 m of the epilimnion (Fig. 1A) was caused by rainfalls during the weeks preceding sampling, since atmospheric precipitation is the only freshwater source to Lake Rogoznica (Žic et al., 2013). Oxygen saturation was at or above 100% in the top 4 m of the epilimnion and steadily decreased until depletion at ∼9 m (Fig. 1A). The chemocline, a sulfide-oxygen interface, was located at 8.5-9 m depth and accompanied by a sharp peak in turbidity (Fig. 1B), indicative of a dense microbial population and the formation of colloidal zero-valent sulfur (S^0^; Kamyshny et al., 2011). S^0^ was detected at 8, 9 and 10 m depth, with a concentration peak of 77 µmol l^-1^ at 9 m (Fig. 1C). Thiosulfate and sulfide were only detected below 8 m, and reached maximal concentrations of 12 µmol l^-1^ and 2.7 mmol l^-1^, respectively (Fig. 1C). DIC concentrations in Lake Rogoznica steadily increased with depth and ranged between 3.4-5.6 mmol l^-1^ (Fig. 1C). These values are at least 1 mmol l^-1^ higher than measured and modeled DIC concentrations in Adriatic surface seawater (e.g. Cossarini et al., 2015; Gemayel et al., 2015). Carbonate mineral (calcite and dolomite) dissolution could have been an important DIC source in Lake Rogoznica, as it has been previously reported for other karst and epikarst hosted surface water bodies and aquifers in Croatia and elsewhere (e.g. Barešić et al., 2011; Florea et al., 2016). Alternatively, organic matter (OM) mineralization in the lake sediment and anoxic water column could have resulted in DIC build-up during water column stratification.

**Figure 1.**
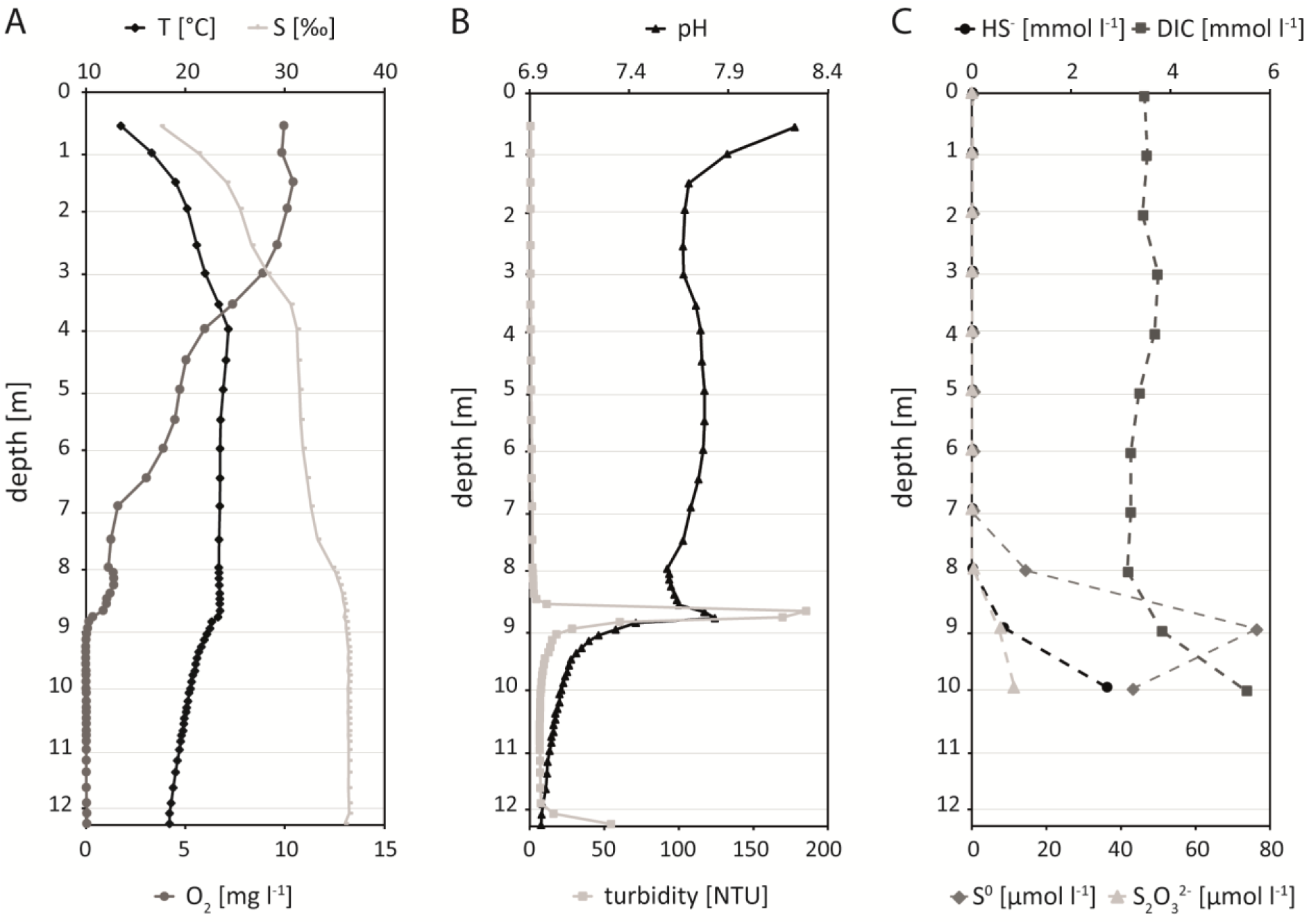
Hydrographical profile and reduced sulfur speciation in a depth profile of Lake Rogoznica collected in April 2015. From left to right: **A** Temperature (T) (°C), salinity (S) (‰) (top x axis), and dissolved oxygen (O_2_) concentration (mg l^-1^) (bottom x axis) profiles. **B** pH (top x axis) and turbidity (NTU) (bottom x axis) profiles. **C** Sulfide (HS^-^) concentration (mmol l^-1^), dissolved inorganic carbon (DIC) concentration (mmol l^-1^) (top x axis), elemental sulfur (S^0^) concentration (µmol l^-1^), and thiosulfate (S_2_O_3_^2-^) concentration (µmol l^-1^) (bottom x axis) profiles.

### Photosynthetic pigment analysis

During stratification, Lake Rogoznica contains a high biomass of anoxygenic photosynthetic bacteria in anoxic waters, while oxygenic phototrophs (i.e. cyanobacteria and algae) inhabit the oxic epilimnion (Malešević et al., 2015; Pjevac et al., 2015). To assess the depth distribution of phototrophic microorganisms, we analyzed the photosynthetic pigment content of a lake water depth profile collected on the day of experimentation. In the upper 8 m of the water column we detected mostly Chl *a*, indicative of the presence of cyanobacteria and algae (oxygenic phototrophs; Scheer, 2006). Chl *a* reached a maximum of 5.0 nmol l^-1^ at 4 m depth, with a secondary peak (4.1 nmol l^-1^) at 8 m depth (Fig. 2). No Chl *a* was detected in samples collected from 9 m and 10 m depth (Fig. 2). In the epilimnion (0-8 m), small amounts of BChl *a* (<5% of identified pigments), likely originating from aerobic anoxygenic species, were detected (Fig. 2). The water samples collected at the chemocline (9 m) and in the hypolimnion (10 m) contained 80-to 110-fold higher concentrations of BChl *a* (11.1-14.5 nmol l^-1^) than the epilimnion, reflecting the presence of GSB, PSB, and potentially other anoxygenic photosynthetic bacteria (Scheer, 2006). No BChl *b, c* or *d* were detected. However, at the chemocline (9 m) and in the hypolimnion (10 m), very high concentrations of BChl *e* (8.1-18.4 nmol l^-1^), only known for low-light adapted, brown-colored GSB (Chew and Bryant, 2007), were detected (Fig. 2). Interestingly, even above the anoxic chemocline (at 8 m), BChl *e* concentrations (0.5 nmol l^-1^) exceeded BChl *a* concentrations (0.1 nmol l^-1^). Since BChl *e* is only found in GSB, while BChl *a* is present in a variety of anoxygenic phototrophic bacteria, the BChl *e* / BChl *a* ratio supports the previously reported dominance of GSB throughout the chemocline of Lake Rogoznica (Pjevac et al., 2015).

**Figure 2.**
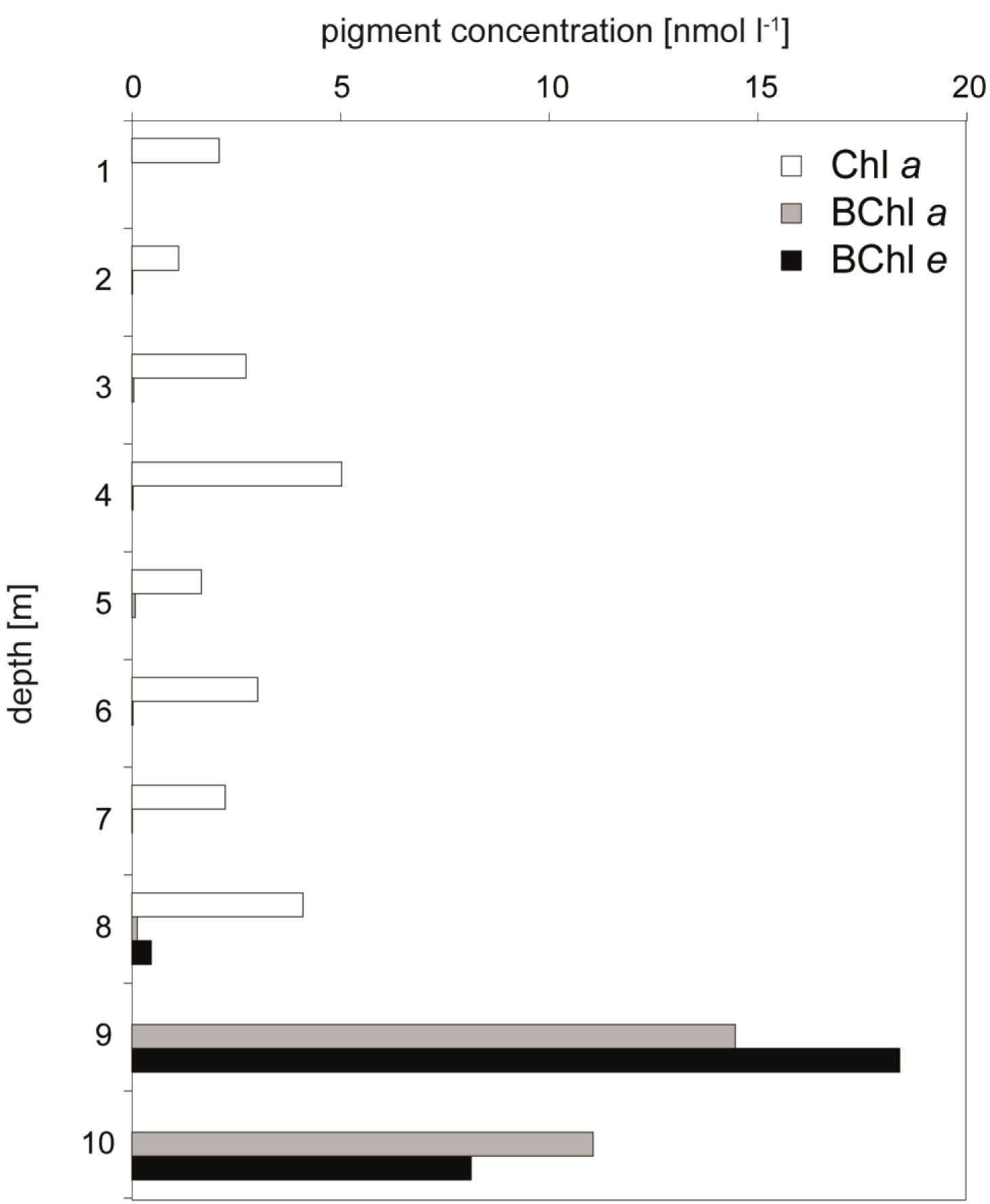
Depth profile of Chlorophyll *a* (Chl *a*), Bacteriochlorophyll *a* (BChl *a*), and Bacteriochlorophyll e (BChl *e*) concentrations (nmol l^-1^) in Lake Rogoznica.

### GSB and PSB in Lake Rogoznica

To determine the community composition of anoxygenic phototrophic bacteria in Lake Rogoznica, we sequenced metagenomes from chemocline (9 m) and hypolimnion (10 m) samples and assessed to relative abundance of GSB and PSB in these samples via 16S rRNA gene sequence read mapping and reconstruction (Fig. S2). In the chemocline (9 m), GSB-related reads accounted for 13% of the total reads mapped to 16S rRNA genes, while PSB-related reads accounted for 5%. In the hypolimnion (10 m) sample, the relative contribution of GSB-and PSB-related reads mapped to 16S rRNA gene sequences were 17% and 2%, respectively (Fig. S2).

From the metagenomic data, we reconstructed three nearly complete high quality metagenome assembled genomes (HQ MAGs) related to anoxygenic phototrophs (Table 1). Two GSB-related HQ MAGs displayed a high average nucleotide identity (>98%) to genomes of validly described *Chlorobiaceae* species (Fig. S3; Table 1). The more abundant GSB in Lake Rogoznica, represented by HQ MAG C10 (Table 1), was identified as a strain of *Chlorobium phaeobacteroides* (Overmann et al., 1992). This is well in line with the high concentration of BChl *e* measured in chemocline and hypolimnion samples, as BChl *e* is the dominant photosynthetic pigment in the low light adapted *Chlorobium phaeobacteroides* strain BS1 (Overmann et al., 1992). The second GSB HQ MAG, B10, was closely related to *Prosthecochloris aestuarii* DSM 271 (Gorlenko, 1970), and represents a strain of this species. Based on read coverage, it was present at a much lower abundance at the time of sampling (Fig. S4). The genomic potential of both HQ MAGs from GSB overlapped with the genomic potential of their close relatives isolated in pure culture: both HQ MAGs show the genomic potential of an obligatory anoxygenic photolithoautotroph.

**Table 1:** Genomic features of recovered metagenome assembled genomes (MAGs) related to anoxygenic phototrophs.

| MAG | Contigs | CDS<sup>1</sup> | Genome size [Mb] | GC content | Completeness | Contamination |
| --- | --- | --- | --- | --- | --- | --- |
| *Halochromatium* sp. (A10) | 43 | 3,913 | 4.19 | 60.8% | 82.9% | 1.8% |
| *Prosthecochloris aestuarii* spp. (B10) | 32 | 2,070 | 2.26 | 50.1% | 92.8% | 0.9% |
| *Chlorobium phaeobacteroides* spp. (C10) | 40 | 2,009 | 2.13 | 44.1% | 91.9% | 0% |

<sup>1</sup>coding domain sequences

For the PSB-related HQ MAG A10 no closely related genome sequence could be identified. However, the two almost identical (1 nucleotide difference) 16S rRNA gene copies encoded in HQ MAG A10 displayed 99% sequence identity to the 16S rRNA gene sequence of *Halochromatium roseum* (Kumar et al., 2007) and the 16S rRNA gene sequence of the PSB species isolated by FACS from Lake Rogoznica in our previous study (Pjevac et al., 2015; Fig. S5). Thus, MAG A10 provides the first genomic insights into the genus *Halochromatium* within the family *Chromatiaceae*. The high 16S rRNA sequence identity of 99% is in the absence of a genome sequence suggestive, yet insufficient for the assignment of this MAG to *Halochromatium roseum.*

All three validly described *Halochromatium* species - *Halochromatium salexigens* (Caumette et al., 1988), *Halochromatium glycolicum* (Caumette et al., 1997), and *Halochromatium roseum* (Kumar et al., 2007) harbor BChl *a* as their main photosynthetic pigment. They require elevated NaCl concentrations (>1%) and grow photolithoautotrophically by oxidation of reduced sulfur compounds. Additionally, the capability to assimilate at least some organic carbon compounds (e.g. acetate, pyruvate, fumarate, succinate, malate, glycolate) during mixo-or heterotrophic growth was demonstrated for all strains. Furthermore, sulfur-dependent chemolithotrophic growth (both autotrophic and heterotrophic) under microaerobic conditions has been demonstrated for *Halochromatium salexigens* and *Halochromatium glycolicum* (Caumette et al., 1988; Caumette et al., 1997; Kumar et al., 2007). In accordance, the here retrieved *Halochromatium* sp. HQ MAG A10 encodes the potential for lithotrophic growth via reduced sulfur compound oxidation and CO_2_ fixation via the Calvin-Benson-Bassham cycle. All genes essential for phototrophic growth and chemotrophic growth with O_2_ as terminal electron acceptor were present (Fig. 3). Moreover, this HQ MAG encodes the capability for N_2_ fixation and urea utilization (Fig. 4) - metabolic features that were not previously reported for members of the genus *Halochromatium*. Finally, the HQ MAG A10 encodes genes for mixo-or heterotrophic metabolism, including glycolate and sucrose utilization, as well as glycogen, polyphosphate, and PHB storage and utilization (Fig. 4).

**Figure 3.**
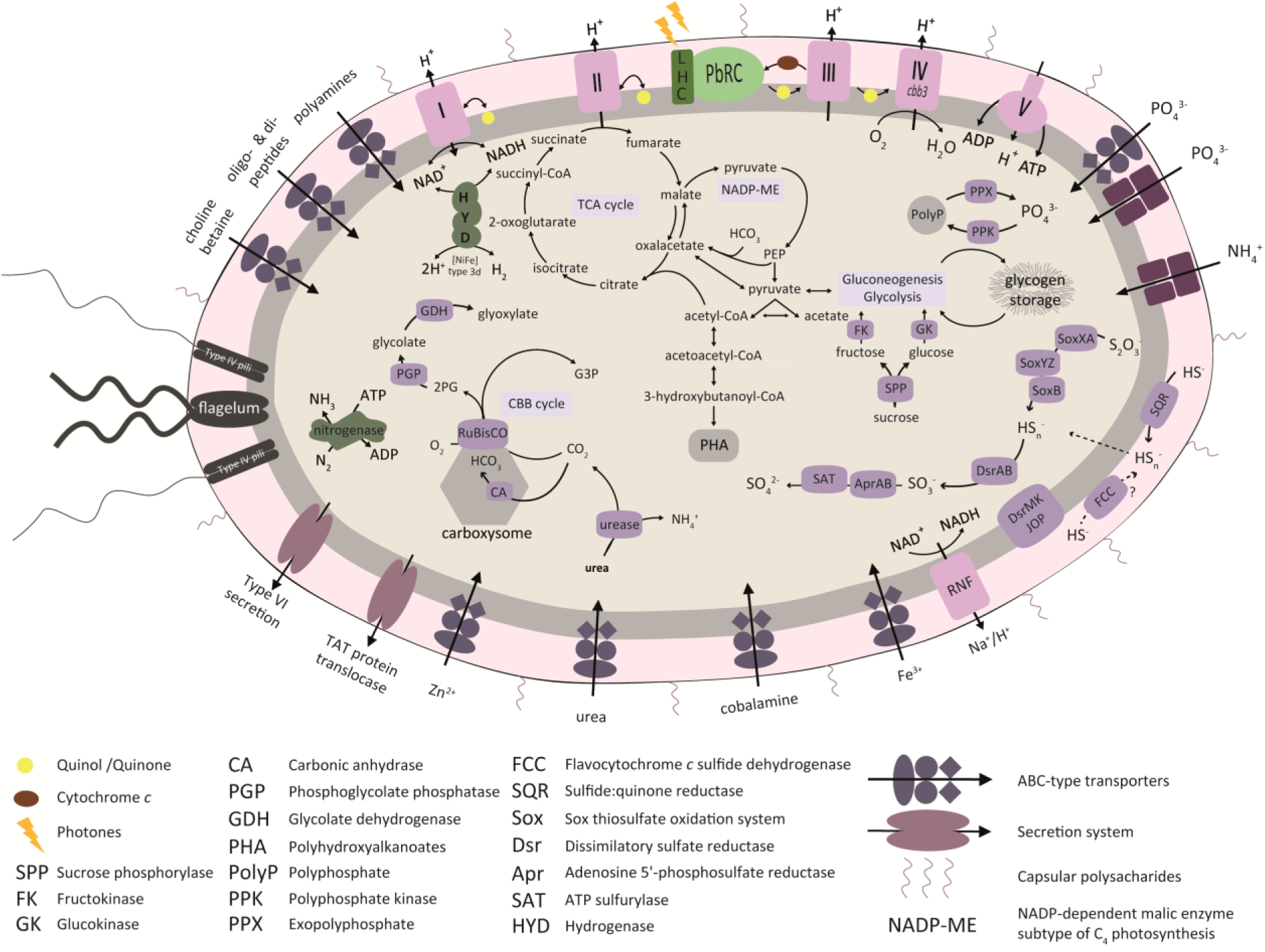
Illustration of a *Halochromatium* sp. cell, depicting select aspects of the metabolic potential inferred from the nearly-complete metagenome assembled genome. Particular focus was directed at the genetic inventory related to photosynthesis, sulfur and nitrogen metabolis m, inorganic and organic carbon utilization, and PHA, glycogen and polyphosphate storage. Respiratory chain membrane complexes are labeled with roman numerals.

**Figure 4.**
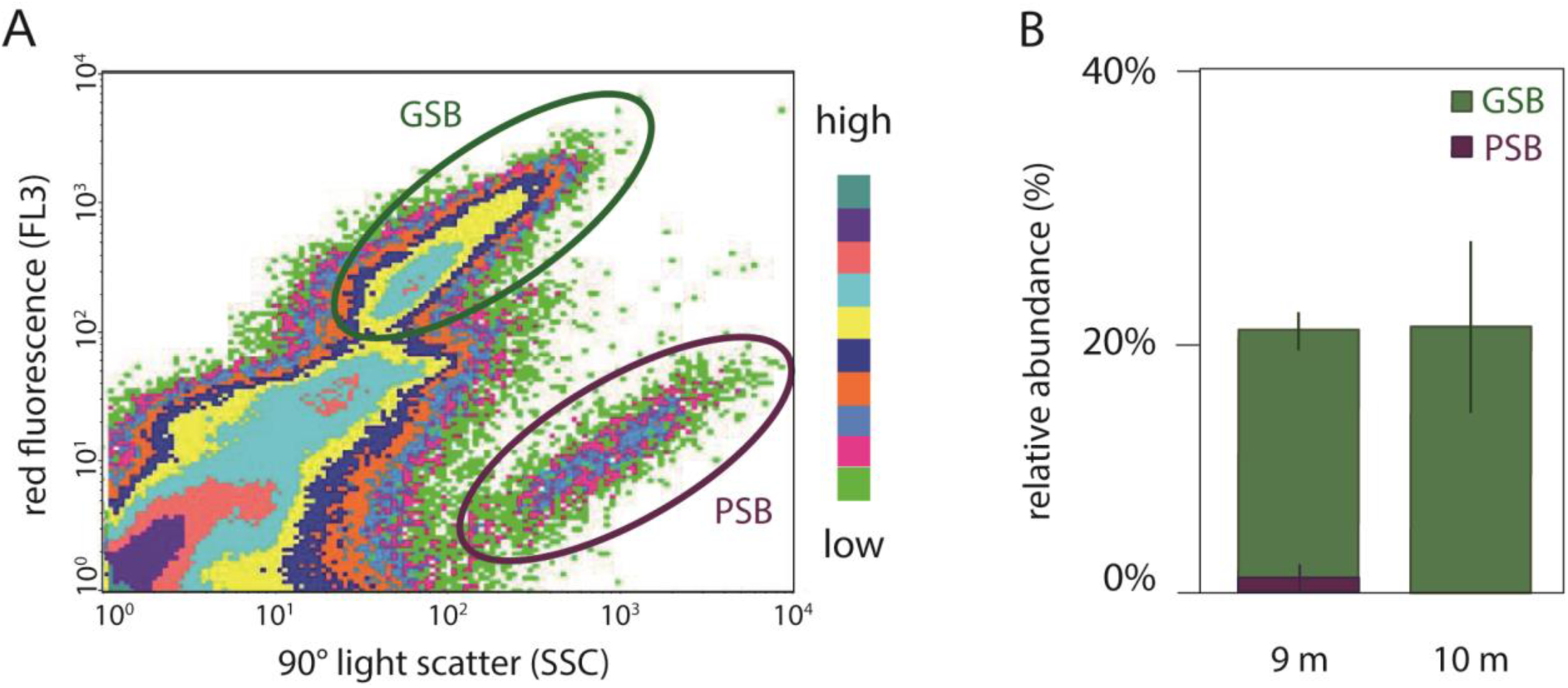
**A** An exemplary sort plot of red fluorescence (FL3; y-axis) versus 90° light scatter (SSC; x-axis) recorded during FACS of a ^14^C-bicarbonate incubated chemocline (9 m) sample. The populations identified as GSB (green circle) and PSB (red circle) are highlighted. The color scale bar indicates event frequencies. **B** Relative abundance of sorted GSB (green) and PSB (purple) populations in the chemocline (9 m) and hypolimnion (10 m) samples.

### FACS sorting and quantification of GSB and PSB populations

To assess the contribution of PSB and GSB to DIC fixation in Lake Rogoznica, we implemented a FACS-based workflow to differentially sort the GSB and PSB populations after *in situ* incubations with radioisotope labeled bicarbonate. Briefly, after sorting and quantification of total microbial cells based on an unspecific DNA stain, the red fluorescence (FL3) of photosynthetic pigments upon excitation with green laser light (488 nm) is used as a diagnostic feature to distinguish phototrophs from other microorganisms. To further distinguish different pigmented populations, both forward scatter (FSC) profiles (reflecting differences in cell size) and 90° side scatter (SSC) profiles (reflecting cell granularity) can be used. In fact, combinations of FL3, FSC and SSC profiles have previously been used to distinguish PSB and GSB from each other and other microorganisms in cultures and environmental samples (Casamayor et al., 2007; Pjevac et al., 2015; Zimmermann et al., 2015). Here, we opted for identification, sorting and quantification of GSB and PSB populations based on FL3 versus SSC profiles, as this yielded the clearest distinction between populations (Fig. 4).

In the chemocline samples, we detected a low abundance (1.0 ±1.0% of all cells) population displaying a FL3 vs. SSC profile resembling that of the PSB previously selectively sorted from Lake Rogoznica (Pjevac et al., 2015). Another, significantly more abundant (18.4 ±1.4%) population was identified on FL3 vs. SSC profiles in the chemocline (9 m) samples (Fig. 4). Taking in account the absence of oxygenic phototrophs at and below the chemocline and the matching relative abundance of GSB in chemocline samples (Fig. S2), we identified this population as GSB (Fig. 4). In the hypolimnion sample (10 m), the abundance of the PSB was too low for sorting, while the GSB accounted for 19.7 ±6.4% of all cells (Fig. 4).

### DIC fixation by anoxygenic phototrophic bacteria in Lake Rogoznica

Previous studies, performed in other aquatic stratified environments, combined *in situ* incubations with isotopically labeled bicarbonate with incubations in the dark or in the presence of oxygenic photosynthesis inhibitors (e.g. 3-(3′,4′-dichlorophenyl)-1,1′-dimethyl urea -DCMU) to distinguish DIC fixation by anoxygenic phototrophs, oxygenic phototrophs and chemolithoautotrophs (e.g. Camacho et al., 2001; Casamayor et al., 2001; García-Cantizano et al., 2005; Marschall et al., 2010; Fontes et al., 2011; Morana et al., 2016). Our approach, on the other hand, requires only *in situ* ^14^C-bicarbonate incubations followed by FACS and scintillography to quantify the contributions of individual anoxygenic phototrophic populations to DIC fixation. Using this method, cell specific rates of *in situ* DIC fixation were determined by scintillography for total cell sorts (i.e. all SYBR Green-I stained cells), as well as GSB and PSB specific cell sorts (i.e. subpopulations of all SYBR Green-I stained cells differentially based on their FL3 vs. SSC profile).

Average *in situ* per cell DIC fixation rates were similar in epilimnion (15.1 ±6.0 amol C cell^-1^ h^-1^) and chemocline (12.4 ±2.8 amol C cell^-1^ h^-1^) samples, while average *in situ* per cell DIC fixation rates in hypolimnion samples (1.2 ±1.2 amol C cell^-1^ h^-1^) were not higher than the background signal measured in dead controls (0.7 ±0.1 amol C cell^-1^ h^-1^) (Fig. 5). Average *in situ* per cell DIC fixation rates of the chemocline-dwelling GSB population from Lake Rogoznica (60.4 ±14.7 amol C cell^-1^ h^-1^; Fig. 5) were in a similar range to rates calculated for the *Chl. clathratiforme* (GSB) cells from Lake Cadagno (Musat et al., 2008). The average *in situ* per cell DIC fixation rates of the chemocline-dwelling PSB population (27.1 ±11.1 amol C cell^-1^ h^-1^; Fig. 5) were in the same order of magnitude as rates determined for chemocline-dwelling GSB in this and other studies (Musat et al., 2008; Storelli et al., 2013). However, these values are up to three orders of magnitude lower than rates determined for PSB populations in Lake Cadagno (Musat et al., 2008; Storelli et al., 2013). Notably, DIC fixation rates of the multiple, co-occurring PSB populations in Lake Cadagno also varied by up to 100-fold (Musat et al., 2008; Storelli et al., 2013).

**Figure 5.**
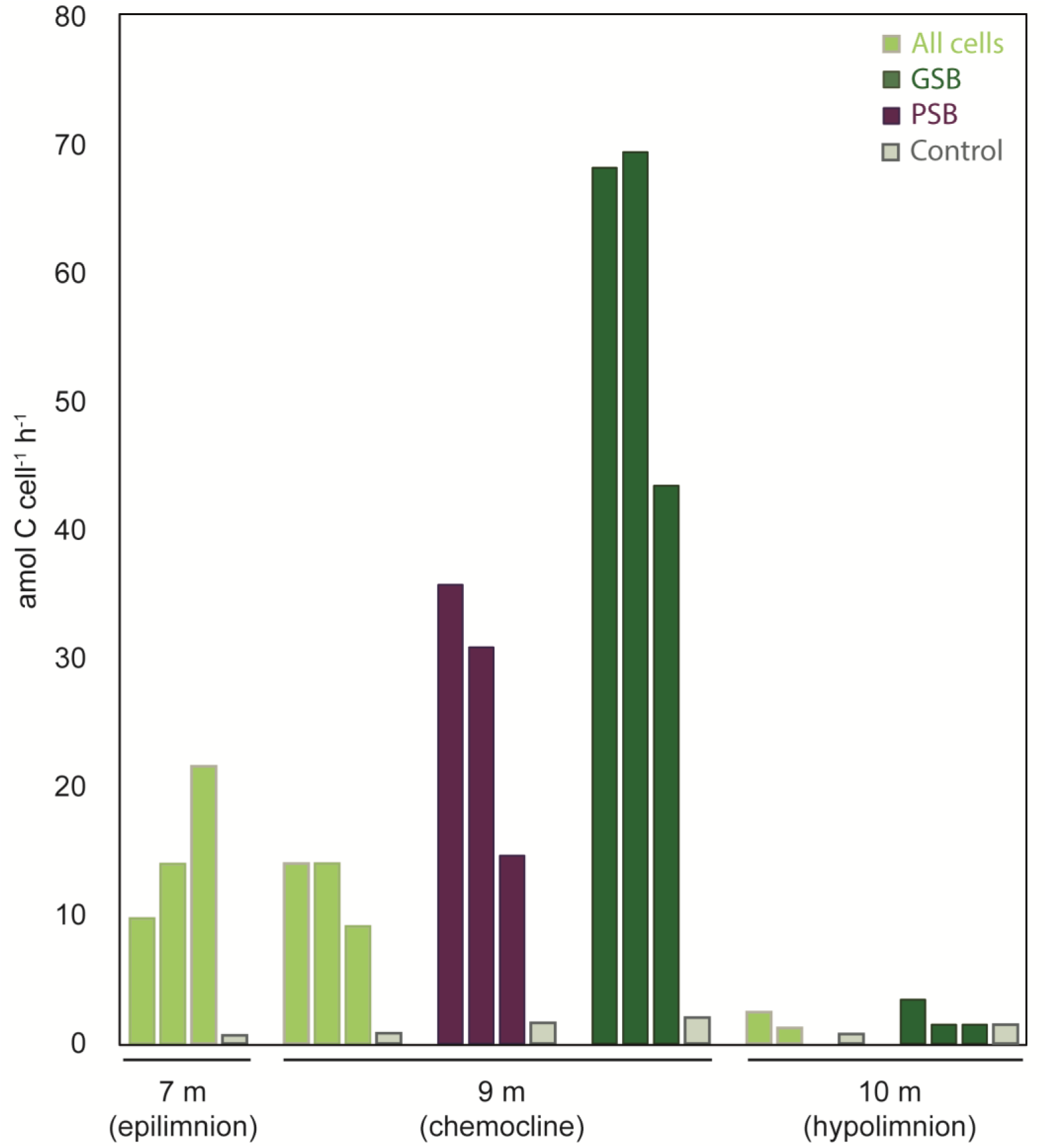
Averaged cell specific carbon assimilation rates (in amol C cell^-1^ h^-1^) plotted separately for each sorted population from epilimnion (7 m), chemocline (9 m) and hypolimnion (10 m) samples. Sorts of all cells (100,000 cells/sort) are shown in light green, sorts of GSB (50,000 cells/sort) are shown in dark green, and sorts of PSB (25,000 cells/sort) are shown in purple. Averaged cell specific carbon assimilation rates (in amol C cell^-1^ h^-1^) for sorts of all cells (100,000 cells/sort) in dead controls are depicted in gray for each depth.

By integrating average cell abundances with average *in situ* per cell DIC fixation rates, we determined that the chemocline-dwelling GSB population was responsible for almost the entire (89.3 ±7.6%) *in situ* DIC fixation measured in this study, whereas the low abundant PSB population contributed only 1.8 ±1.4%. This is in contrast to the disproportionally high contribution of PSB to *in situ* DIC fixation reported for the chemocline of Lake Cadagno (Musat et al., 2008; Storelli et al., 2013).

Finally, the hypolimnion-dwelling GSB population (2.1 ±1.1 amol C cell^-1^ h^-1^) in Lake Rogoznica did not assimilate significant amounts of labeled DIC (Fig. 5). This result is consistent with previously observed low DIC uptake by anoxygenic phototrophs in the hypolimnion (e.g. Guerrero et al., 1985; Camacho and Vicente 1998) and can be attributed to low light quality through self-shading.

## Conclusions

We show that anoxygenic photosynthesis-driven DIC fixation in Lake Rogoznica in spring 2015 was primarily mediated by a highly abundant chemocline-dwelling GSB population, whereas the co-occurring PSB contributed only marginally to DIC fixation. Our radioisotope-labeling and FACS-based workflow facilitated unprecedented insights into DIC fixation by environmental populations of anoxygenic phototrophs. Our results contrast previous findings from Lake Cadagno and illustrate that high contributions of rare PSB populations to DIC fixations cannot be generalized for other stratified lakes. Further research is needed to obtain a more representative and comprehensive picture of PSB vs. GSB mediated DIC fixation in environments inhibited by anoxygenic phototrophs. Furthermore, the here presented workflow can be readily adapted for analyze various co-occurring phototrophic microbial populations, and also used to investigate other aspects of the *in situ* metabolism of physiologically flexible PSB. For example, photoheterotrophic activity, a prominent metabolic feature encoded in the here recovered *Halochromatium* MAG and other PSB genomes (Berg et al., 2019; Luedin et al., 2019), could be quantified by tracking the incorporation of radiolabeled organic substrates into environmental PSB populations.

## Experimental procedures

### Study site and sample collection

Lake Rogoznica (Fig. S1) is a small (∼1 ha, 15 m max depth) seawater lake on the Dalmatian peninsula Gradina, Croatia (43°32′N, 15°58′E). It is located at approximately 100 m distance from the seashore, exchanging water with the Adriatic Sea through subsurface channels in the porous limestone characteristic to Croatian coastal areas (e.g. Žic et al., 2013). The lake water is rich in nutrients and dissolved organic carbon (DOC). The hypolimnion is highly sulfidic during stratification (Pjevac et al., 2015). On April 2^nd^, 2015 hydrographical parameters (dissolved oxygen concentration [O_2_], temperature [T], conductivity [Sm], turbidity [t] and pH) of a vertical lake profile were recorded with a Hydrolab DS5 multiparameter water quality probe (OTT Hydromet, Germany). Water samples for DIC and reduced sulfur species (RSS) analyses were collected with a vertically lowered 5-liter Niskin bottle (General Oceanics, USA) in 1 meter intervals between the surface [∼10-60 cm] and 10 m water depth. At 7 m (epilimnion), 9 m (chemocline) and 10 m (hypolimnion) below surface, larger sample volumes were collected for activity incubations and metagenome sequencing.

### Sample preservation and chemical analyses

Samples for DIC analyses were collected in 1.5 ml amber borosilicate vials (Zinsser Analytic Qualyvials); 15 µl saturated mercury chloride solution (2.7 mol l^-1^) were added and vials were closed with polytetrafluoroethylene (PTFE) coated septa and screw caps. DIC was measured by infrared spectroscopy (Analytik Jena multi N/C 2100s) as CO_2_ liberated from the original sample in a 10% phosphoric acid trap. Samples for reduced sulfur species analyses were collected in 1.5 ml amber borosilicate vials (Zinsser Analytic Qualyvials) and preserved with ZnCl_2_ (final concentration 4%). Sulfide concentrations were determined spectrophotometrically in ZnCl_2_ fixed samples as described before (Cline, 1969). Zero-valent sulfur was converted to thiosulfate via sulfitolysis (Jørgensen et al., 1979; Ferdelman et al., 1991). Briefly, ZnCl_2_ fixed samples were buffered with HEPES (pH 8) and reacted with a 2% sulfite solution at 70°C for 12 h to yield thiosulfate. Parallel samples with no added sulfite were used as a control for background thiosulfate concentrations. The thiosulfate products were then derivatized with monobromobimane as described in Zopfi et al., (2004). Derivatized samples were analyzed on an Acquity H-class UPLC system (Waters Corporation, USA) equipped with an Acquity UPLC BEH C8 column and controlled with the Empower III software. Sample temperature was maintained at 4°C, column temperature was 40°C, and the sample injection volume used was 1 µl. The mobile phase consisted of acetic acid (0.25 v/v), pH 3.5 (A), and 100% UPLC-grade methanol (B). The following gradient conditions were used at a constant flow rate of 0.65 ml min^-1^: start, 5% B; 0-4 min, ramp of curve 6 (linear) to 10% B; 4-5 min, ramp of curve 6 to 95% B; 5-6 min, 95% B; 6.01 min, 5% B; 8 min, 5% B; injection of the next sample. The fluorescence detector was set to an excitation wavelength of 380 nm and an emission wavelength of 480 nm.

### Pigment analysis

One l (0-7 m) or 1.5 l (8-10 m) water samples were collected onto 47 mm glass fiber filters (type GF/F, nominal pore size 0.7 µm) using low vacuum. The filters were folded, gently dried using a paper towel, and stored at −80°C. In the laboratory, the filters were homogenized in 7 ml acetone: methanol mixture (7:2) using a glass Teflon tissue homogenizer. The filter debris was removed by centrifugation (10,000 g). Pure extracts were analyzed using the Prominence - *i* LC-2030C HPLC system (Shimadzu, Japan). Pigments were separated on a heated (40°C) Luna 3μC8(2) 100 Å column (Phenomenex Inc., USA) with binary solvent system A: 20% 28 mmol l^-1^ ammonium acetate + 80% methanol, B: 100% methanol. Chlorophyll (Chl) *a*, BChl *a*, and BChl *e* were detected at 665 nm, 770 nm, and 655 nm, respectively. The HPLC system was calibrated using 100% methanol extracts of *Synechocystis* sp. PCC6803, *Rhodobacter sphaeroides* and *Chlorobium phaeobacteroides* with known concentrations of Chl *a*, BChl *a*, and BChl *e*, respectively.

### DNA extraction and metagenome sequencing

DNA for metagenome sequencing was extracted from 6 polycarbonate membrane filters (type GTTP; 0.2 µm pore size, Whatman, UK) per depth (9 m and 10 m) by a phenol/chloroform-based protocol (Massana et al., 1997). Briefly, filters were incubated with lysozyme (1 g l^-1^) at 37°C for 45 min, and proteinase K (0.2 g l^-1^) and SDS (1%) at 55°C for 1 h. Extraction was performed twice with 750 μl of phenol: chloroform (CHCl_3_): isoamyl alcohol (IAA) (25: 24: 1, pH 8) solution and once with 750 μl of CHCl_3_: IAA (24: 1) solution. The aqueous phases of all extractions for each depth were collected and pooled. Finally, 1/10 volume of sodium acetate was added to the pooled aqueous phase and DNA was precipitated for 30 min at −20°C with 1 ml of isopropanol. After a centrifugation step (20 min at 4°C and 20,000 g) the DNA pellet was washed with 500 μl of 70% ethanol. After a second centrifugation step (5 min at 4°C and 20,000 g) the pellet was dissolved in 80 μl of deionized (MQ) water and stored at −20°C until further processing. Aliquots of the DNA extracts were sent to the Max Planck Genome Centre (MP-GC, Cologne) for paired end library preparation and metagenome sequencing on the Illumina HiSeq 2500. Paired end metagenome reads were quality trimmed at a phred score of 15 using the bbduk function of the BBMap package (v. 35.82; https://sourceforge.net/projects/bbmap/). Small subunit (SSU) rRNA gene sequences were reconstructed from the quality trimmed metagenomic reads and classified against the SILVA SSU rRNA gene database using PhyloFlash (v.3.0; Gruber-Vodicka et al., 2019). A metagenome co-assembly of both samples was performed with Spades v. 3.11.1 (Bankevich et al., 2012) and binned with Metwatt v. 3.5.3 (Strous et al., 2012). Three HQ MAGs classified as anoxygenic phototrophs were identified and polished by iterative re-assembly for 10 iterations as described in Mußmann et al., 2017. The phylogenetic affiliation, completeness and redundancy of the HQ MAGs was assessed with MIGA (Rodriguez-R et al., 2018) and the HQ MAGs were automatically annotated in RAST (Aziz et al., 2008). For phylogenetic placement of the *Halochromatium* sp. A10 HQ MAG, the 16S rRNA gene sequences encoded in HQ MAG A10, alongside with a selection of 16S rRNA gene sequences from *Chromatiaceae* isolates and environmental samples were aligned against the SILVA r132 SSU database using the SINA aligner (Pruesse et al., 2012). The alignment was uploaded to the IQ-TREE webserver (Trifinopoulos et al., 2016) for phylogenetic tree reconstruction using default parameters, and the resulting tree was visualized in iTOL (Letunic and Bork, 2016).

### Nucleotide Accession Numbers

Metagenome assembled genomes (MAGs) are available under NCBI genome accessions/RAST project IDs POWB00000000/2049430.3 (*Halochromatium sp.* A10), POWC00000000/290513.3 (*Prosthecochloris sp.* B10) and POWD00000000/290513.4 (*Chlorobium sp*. C10), respectively, while the metagenome raw reads can be retrieved from ENA under the study accession number PRJEB26778.

### Carbon assimilation experiments

Samples for *in situ* ^14^C-bicarbonate assimilation experiments were collected with a 5-liter Niskin bottle at 7 m, 9 m and 10 m water depth as described above. Sample aliquots were transferred to 6 ml exetainer screw cap vials (Labco Limited) directly from the Niskin bottles. A sterile rubber tube and a slow flow rate were used during transfer to avoid oxygen intrusion to anoxic chemocline and hypolimnion samples. Four exetainer vials were overflown and filled at each depth. From one exetainer per depth, 325 µl sample were removed with a syringe through the screw cap septum, and replaced with 325 µl 37% formaldehyde solution, resulting in a final formaldehyde concentration of 2%. Thereafter, 0.10 mmol l^-1^ (7 m), 0.12 mmol l^-1^ (9 m) or 0.15 mmol l^-1^ (10 m) ^14^C-labeled sodium bicarbonate (specific activity 56 mCi mmol^-1^) was added to all 4 exetainers of each depth sample with a glass syringe. The sample-filled exetainers were mounted vertical to carrier rings, and lowered to *in situ* depths of 7 m, 9 m or 10 m, respectively. Incubations were performed at ambient light and temperature conditions between 8 am and 2 pm, to assure light availability for phototrophic activity. After 6 h of incubation samples were retrieved, fixed with 2% formaldehyde (final concentration) as described above, and kept at 4°C until further processing.

### FACS and scintillation counting of sorted cells

Prior to flow cytometry, cells were stained with SYBR Green I (Marie et al., 1997) and large suspended particles were removed by filtration through 5 µm pore-size filters (Sartorius) to avoid clogging of the flow cytometer. Flow sorting was performed using a BD FACSCalibur flow cytometer equipped with a cell sorter and a 15 mW argon ion laser exciting at 488 nm (Becton Dickinson, UK). Autoclaved Milli-Q water was used as sheath fluid. Cell sorting was done at a low flow rate of 12 ± 3 µl min^-1^ or a medium flow rate of 35 ± 5 µl min^-1^ with single cell sort mode to obtain highest purity. The event rate was adjusted with a fluorescence threshold and sorting was performed on a rate of approximately 25-250 particles s^-1^. SYBR Green I stained cells were identified on scatter dot plots of green fluorescence (filter FL1 530/30) versus 90° light scatter. Bacteriochlorophyll fluorescence was identified on scatter dot plots of red fluorescence (filter FL3 650LP) versus 90° light scatter. For subsequent measurements 25,000-100,000 cells were sorted and filtered onto 0.2 µm polycarbonate filters (GTTP, Millipore). Collected cell batches on polycarbonate filters were directly transferred into 6 ml scintillation vials and mixed with 5 ml UltimaGold XR (Perkin Elmer) scintillation cocktail. Radioactivity of sorted cell batches was measured in a liquid scintillation counter (Tri-Carb 2900, Perkin Elmer). The abundance of PSB and GSB in ^14^C-carbon assimilation experiments is given as relative percentage of gated cells identified as PSB/GSB out of all SYBR green-stained cells concurrently counted using flow cytometry.

## Acknowledgements

We thank Iva Soža, Klara Filek, Neven Cukrov and the scientists of IRB research station Martinska for support during sample collection. We are especially grateful to Jasmine S. Berg for determining reduced sulfur species concentrations, and to Cedric Garnier for providing us with the Hydrolab DS5 multiparameter water quality probe. Alastair Gardiner is acknowledged for language editing and Martina Hanusova for preparing the pigment extracts. Financial support was provided by the Max Planck Gesellschaft (MPG), and the Croatian Science Foundation through the BABAS project.

