## Supplementary material for "*In situ* abundance and carbon fixation activity of distinct anoxygenic phototrophs in the stratified seawater lake Rogoznica": Fig. S

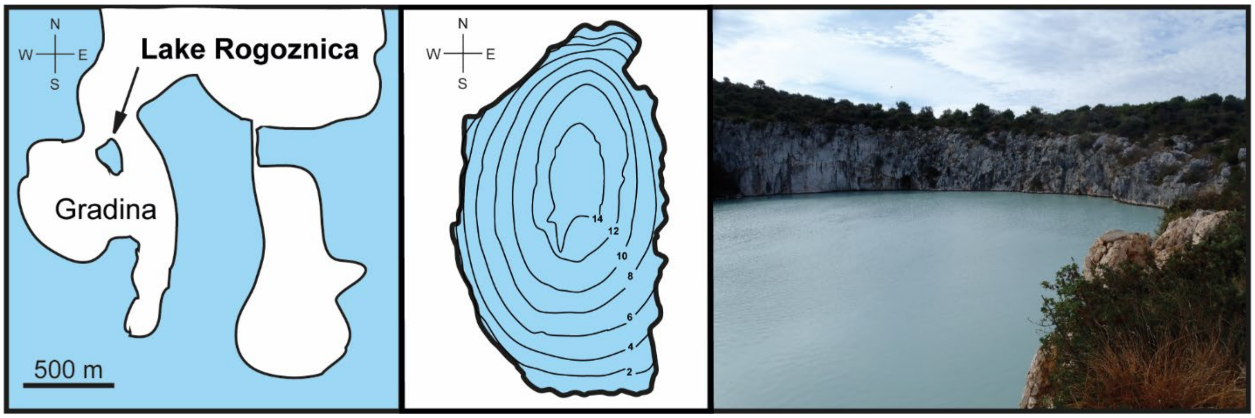


**Figure S1.**

Left panel: location of Lake Rogoznica on the Gradina peninsula. Middle panel: bathymetric map of Lake Rogoznica, depth given in meters. Right panel: photography of Lake Rogoznica from the south-eastern lake shore.

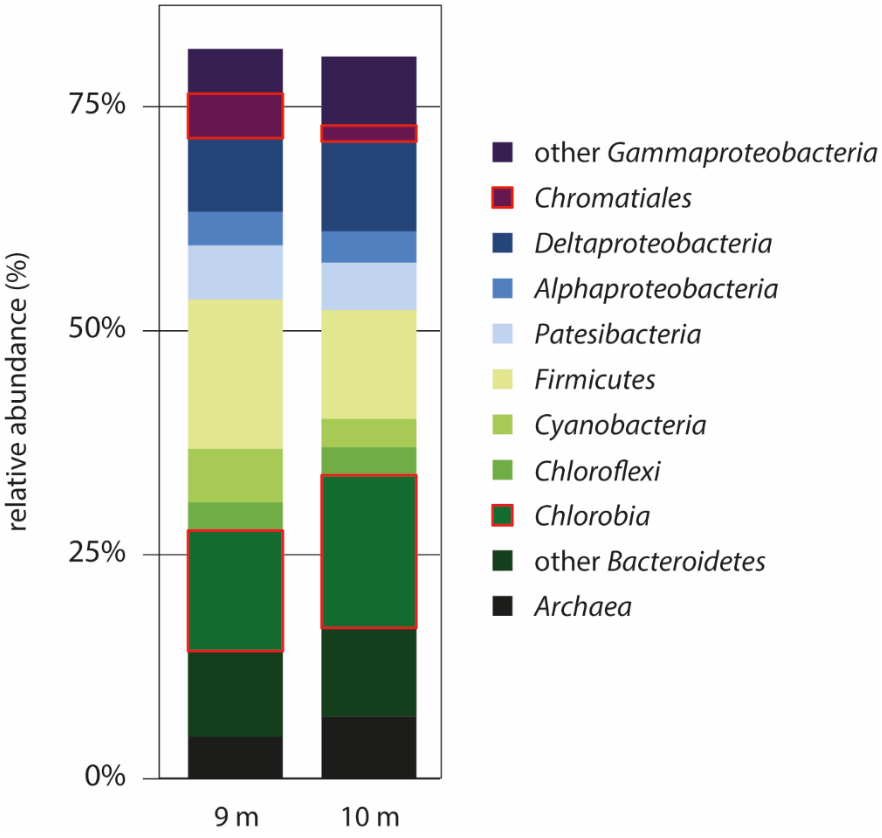


**Figure S2.**

Relative abundance of reads from the chemocline (9 m) and hypolimnion (10 m) metagenomes mapped to 16S rRNA gene sequences, clustered to represent major taxonomic groups in the sample. Sequences related to the order *Chromatiales* and the class *Chlorobia*, harboring the anoxygenic phototrophs occurring in Lake Rogoznica, are highlighted in red.

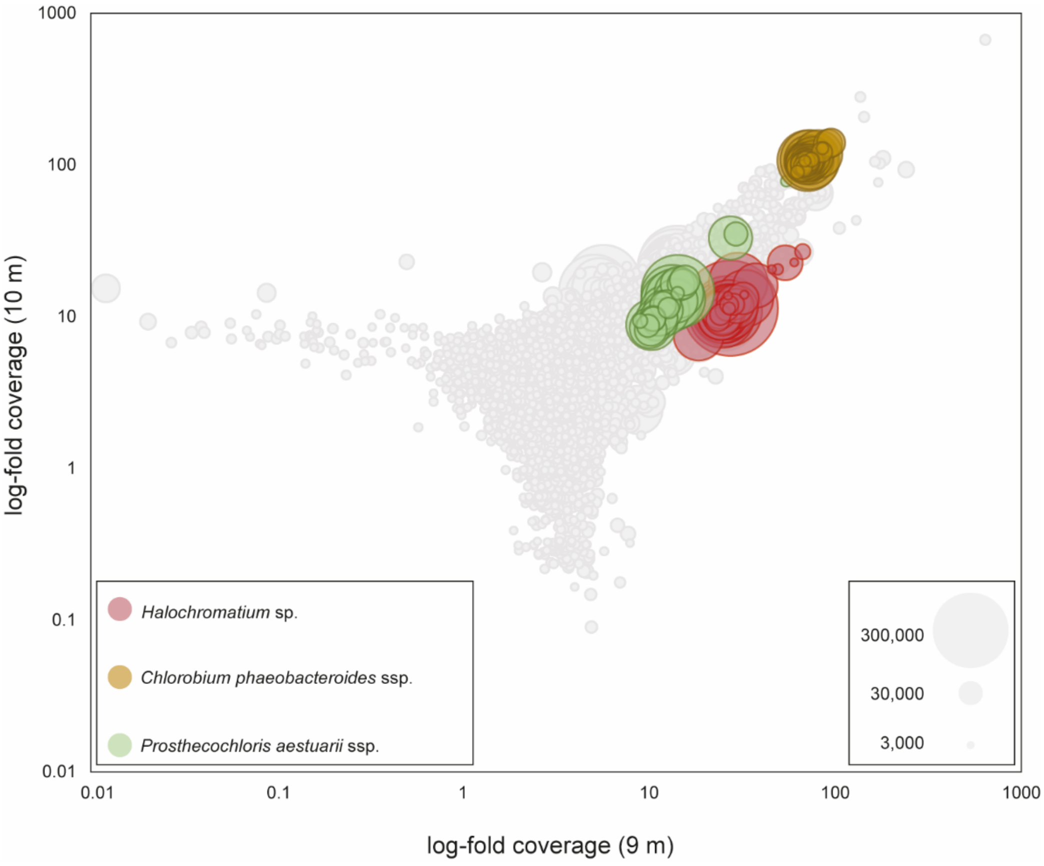


**Figure S3.**

Contigs (> 1,000 bp, min 0.01x/ max 1000x coverage) obtained from a combined bulk assembly of Lake Rogoznica chemocline and hypolimnion samples. The size of circles represents contig size. Larger clusters and clouds of circles indicate contigs with similar coverage. Contours of the three MAGs affiliated to anoxygenic phototrophs are highlighted in the respective colors.

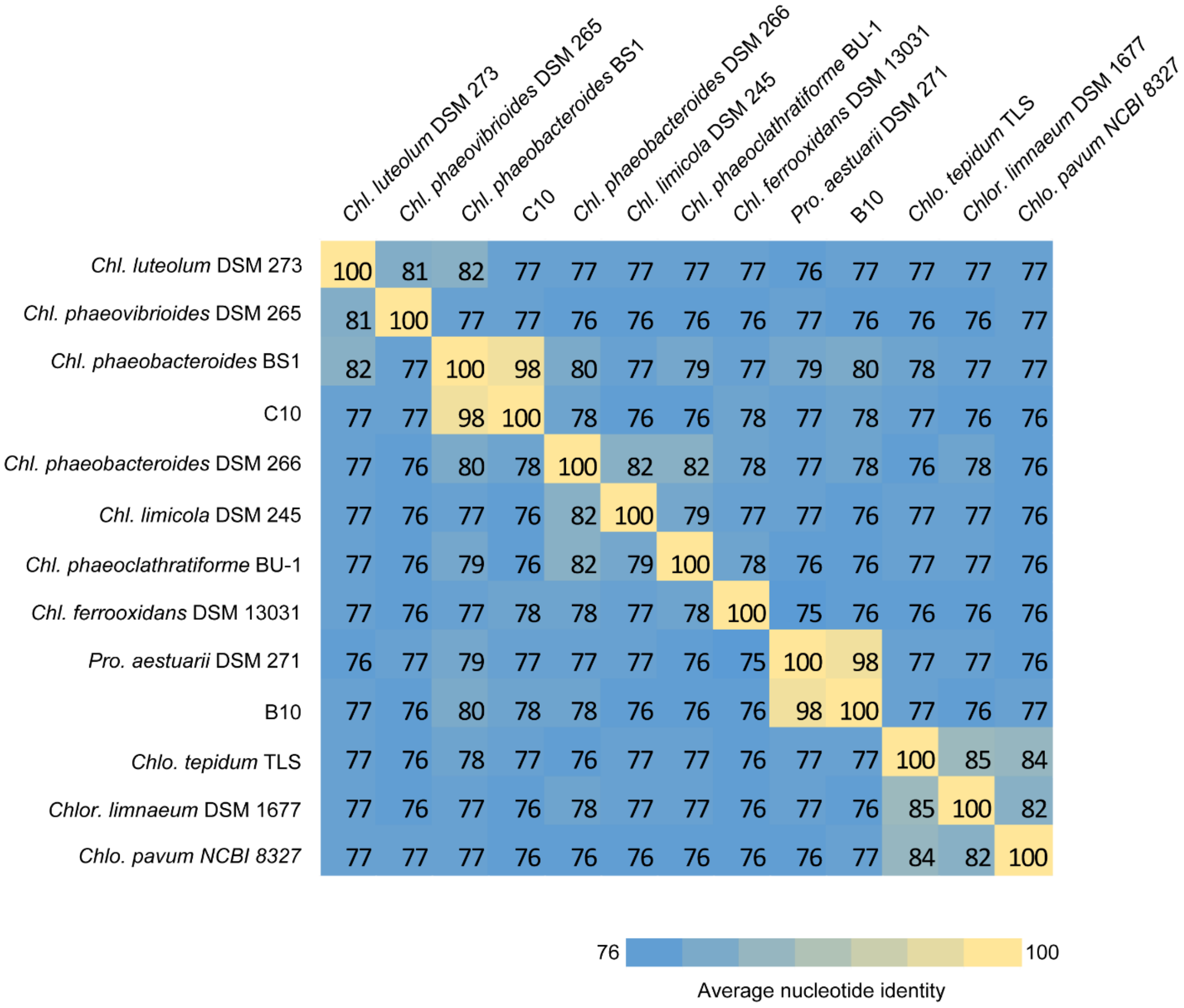


**Figure S4.**

Heatmap depicting average nucleotide identity (ANI) clustering across the *Chlorobiaceae*-related metagenome assembled genomes (MAGs) obtained from the Lake Rogoznica metagenome (C10 and B10) and genomes of other validly described *Chlorobiaceae* species. The color scale bar indicates percent identity.

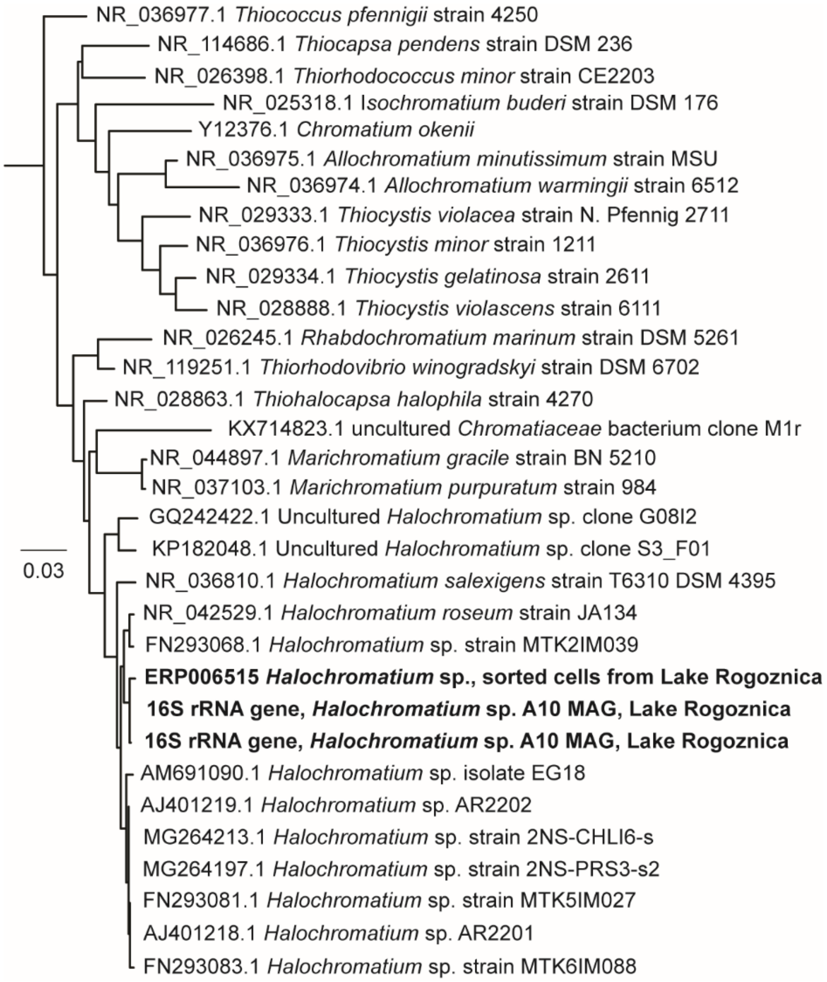


**Figure S5.**

Phylogenetic tree illustration the relation of the two 16S rRNA gene sequences obtained from the *Halochromatium* sp. metagenome assembled genome (in bold) to 16S rRNA gene sequences of isolates from the genus *Halochromatium* and other purple sulfur bacteria isolates (in cursive). The scale bar indicates a sequence dissimilarity of 3%. GenBank accession number are given for each sequence.
